# Epigenetic activation of the prostaglandin receptor EP4 promotes resistance to endocrine therapy for breast cancer

**DOI:** 10.1101/056770

**Authors:** Jeffrey F. Hiken, James I. McDonald, Keith F. Decker, Cesar Sanchez, Jeremy Hoog, Nathan D. VanderKraats, Kyle L. Jung, Margaret Akinhanmi, Matthew J. Ellis, John R. Edwards

## Abstract

Approximately 75% of breast cancers express estrogen receptor α (ERα) and depend on estrogen signals for continued growth. Aromatase inhibitors (AIs) prevent estrogen production and inhibit estrogen receptor signaling, resulting in decreased cancer recurrence and mortality. Advanced tumors treated with AIs almost always develop resistance to these drugs via the up-regulation of alternative growth signals. The mechanisms that drive this resistance—especially epigenetic events that alter gene expression—are however not well understood. Genome-wide DNA methylation and expression analysis of cell line models of acquired aromatase inhibitor resistance indicated that prostaglandin E_2_ receptor 4 *(PTGER4)* is up-regulated after demethylation in resistant cells. Knockdown and inhibitor studies demonstrate that *PTGER4* is essential for estrogen independent growth. Analysis of downstream signaling indicates that *PTGER4* likely promotes AI resistance via ligand independent activation of the ERα-cofactor CARM1. We believe that we have discovered a novel epigenetic mechanism for altering cell signaling and acquiring endocrine therapy resistance. Our findings indicate that *PTGER4* is a potential drug target in AI resistant cancers. Additionally, the epigenetic component of *PTGER4* regulation suggests that further study of *PTGER4* may yield valuable insights into how DNA methylation-targeted diagnoses and treatments can improve AI resistant breast cancer treatment.

## Introduction

Estrogen receptor α(ERα) is necessary for normal human breast development due to its role in processing estrogen growth signals. About 75% of breast tumors express and depend upon ERα^1-3^. Endocrine therapies that disrupt estrogen-ERα binding have a long record of clinical efficacy. Endocrine therapies can be divided into two broad classes: 1) selective estrogen receptor modulators (SERMs) that competitively inhibit estrogen binding to ERα, and 2) aromatase inhibitors (AIs) that prevent the synthesis of estrogen. AIs decrease recurrence and mortality in post-menopausal women when compared to SERMs, such as tamoxifen^4^. Despite this increased efficacy, resistance to AI therapy occurs in nearly all of the advanced or metastatic tumors often through the activation of ligand-independent ER signaling^5^.

Genetic analysis of both patient tumors and cell line models have revealed several mechanisms of acquired AI resistance^5^. Genomic altERαtions such as amplifications, mutations, or translocations can activate *ESR1* in low estrogen conditions^6,7^. Ligand-independent ERα activation can also occur through activation of the PI3K and MAPK signaling pathways at the cell membrane^8^. Activating mutations in the PI3K and MAPK pathways are frequently found in ERα-positive breast cancers^9^. MAPK signaling required for estrogen-independent growth can also be activated by upstream factors such as silencing of the cyclin-dependent kinase CDK10^10^. The downstream effectors of these pathways are responsible for phosphorylation of ERα, which activates it in the absence of estrogen ^11,12^.

Despite improved understanding of potential genetic mechanisms leading to acquired AI resistance, potential epigenetic mechanisms of resistance are not well explored. Almost all cancers exhibit altered DNA methylation, an epigenetic mark that contributes to cancer development^13^ and progression^14^. Epigenetic studies of endocrine therapy resistance have mostly focused on the direct silencing of *ESR1* mediated by either DNA methylation or histone deacetylation^16-23^. However, less is known about how epigenetic changes might contribute to the regulation of transcriptional networks in the development of acquired AI resistance.

In this work, we hypothesized that changes in DNA methylation contribute to acquired endocrine therapy resistance. Resistance to estrogen withdrawal was modeled in ERα-positive cancer cell lines that have been subjected to long-term estrogen deprivation (LTED). LTED cell line models have facilitated the identification of mechanisms of acquired endocrine therapy resistance including increased ERα expression^7^ as well as increased signaling through PI3K, AKT, and MAPK^24-26^. They also have been used to show that PI3K pathway inhibitors induce cell death in ERα-positive cell lines with oncogenic PI3K mutations, suggesting that targeting the PI3K pathway may improve treatment options for a subset of women ^24, 25^.

Genome-wide methylation and expression analysis of LTED cells identified hypomethylation correlated with increased expression of the prostaglandin E_2_ receptor 4 gene *(PTGER4)* in LTED cells. EP4, the *PTGER4* gene product, is a G-protein coupled receptor that activates adenylyl cyclase (AC) and protein kinase A (PKA) in response to prostaglandin E_2_^27^. We find that EP4 activity is necessary for the proliferation of LTED cells. We also show that EP4 up-regulation likely exerts its proliferative effect through PKA-mediated activation of CARM1, which in turn acts as a co-activator with ERα to promote ligand-independent activation of ER-response genes. The significance of these molecular studies elucidating how EP4 is required for estrogen-independent growth was further demonstrated in the identification of *PTGER4* up-regulation in AI-resistance breast tumor samples. The loss of methylation and activation of *PTGER4* represents a possible mechanism of acquired endocrine therapy resistance that can be therapeutically targeted.

## Results

### DNA methylation is altered in a model of acquired resistance to endocrine therapy

To understand potential epigenetic causes of acquired resistance to endocrine therapy, we performed genome-wide methylation and transcriptome analysis in the MCF7 cells conditioned to grow in the absence of estrogen (MCF7-LTED, long-term estrogen deprived) (GSE45337, GSE74943; GSE45337 is public, reviewer link for GSE74943:http://www.ncbi.nlm.nih.gov/geo/query/acc.cgi?token=ubihoogupisliah&acc=GSE74943. Since MCF7 cells do not express aromatase (as verified by our RNA-seq data), aromatase inhibitor resistance is modeled by estrogen withdrawal. After the removal of estrogen, most cells die; however, a few survive and eventually begin to proliferate in the absence of estrogen (Fig. 1a)^24^.

**Figure 1.**
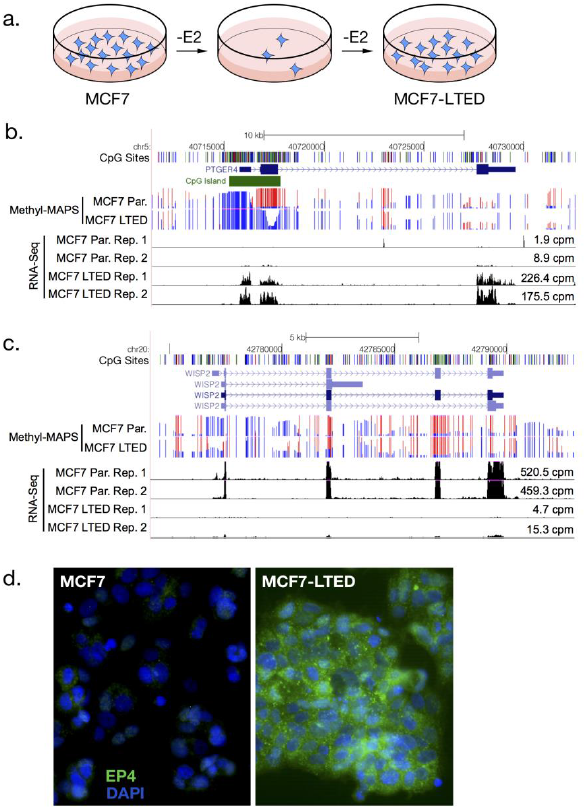
Genome-wide methylation and expression analysis MCF7-LTED cells. (a) Flowchart showing the production of the MCF7-LTED cell line via removal of estrogen (E2) from the growth media. Proliferating cells after estrogen deprivation comprise the MCF7-LTED cell line. (b and c) Genome browser view of Methyl-MAPS methylation and RNA-seq expression data for (b) PTGER4, the gene that encodes EP4, and (c) WISP2. Red and blue lines indicate coverage of methylated and unmethylated fragments, respectively. Individual CpG sites are noted by tics in black at the top track. (d) Immunofluorescence staining of EP4 (green) and DAPI (blue) in MCF7 and MCF7-LTED cells.

Genome-wide methylation analysis using Methyl-MAPS^28^ indicated genome-wide hypomethylation in MCF7-LTED compared to MCF7 cells with 245 644 CpG sites losing methylation and 28 751 sites gaining methylation. Analysis of these sites indicates that the majority of these changes occurred in the transposable elements (Supplementary Fig. 1a). Previously, it was shown that hypomethylation induced by 5-azacytidine increases estrogen-independent growth^29^, which suggests a general mechanism whereby loss of methylation in breast tumors could contribute to estrogen-independent growth and thus endocrine therapy resistance.

### LTED Cells Up-regulate ERα response genes and Potential Resistance Genes

RNA-seq analysis indicated 443 up-and 353 down-regulated genes in MCF7-LTED cells relative to MCF7. We searched the promoters of up-and down-regulated genes for changes in methylation from 500 bp upstream to 1 kb downstream of the transcription start sites, since these regions often correlate with gene expression changes^30,31^. We identified seven genes with promoter methylation changes that associated with expression changes. Identified genes included *PTGER4* and *WISP2* (Supplementary table 1). We validated correlations by examining whether differentially methylated regions correlated with expression across 632 breast tumors and 98 normal breast tissue samples from The Cancer Genome Atlas (TCGA, Supplementary Fig. S4). *WISP2* showed a marked increase in methylation 3’ of the TSS, which was accompanied by a dramatic decrease in expression (Fig. 1c). It was previously shown that knockdown of *WISP2*, a WNT-signaling interacting protein, promoted estrogen-independent growth in MCF7 cells^32^. WNT-signaling has been frequently implicated in epithelial-mesenchymal transition (EMT), and ontology pathway analysis indicated that genes up-regulated in MCF7-LTED cells were enriched for TGF-beta dependent induction of EMT. However, visual inspection of MCF7-LTED cells did not indicate an obvious EMT phenotype and loss of E-cadherin expression was not observed based on RNA-seq data.

Instead, we focused on the *PTGER4* gene that encodes the EP4 receptor. We chose EP4 because it showed the largest expression increase of the candidates, showed a strong and clear change in methylation, and PTGER4 presents a potential druggable target. EP4 is a G-protein coupled receptor that activates AC and PKA in response to prostaglandin E_2_ (Yokoyama et al., 2013). The *PTGER4* gene has a CpG island associated with its promoter that lost methylation in the transition from MCF7 to MCF7-LTED cells (Fig. 1b). Methylation changes were validated using Methyl-Screen (Supplementary Fig. S2a)^34^. This loss of methylation is accompanied with a greater than 20-fold gain in *PTGER4* expression as indicated by RNA-seq (Fig. 1b, Parental cpm = 1.9, 8.9; LTED cpm; = 226.4, 175.5) and confirmed by RT-qPCR (Supplementary Fig. S2b). Furthermore, MCF7 cells treated with 5-azacytidine expressed higher levels of *PTGER4*^33^, consistent with the hypothesis that DNA hypomethylation directly regulates expression. Increased mRNA levels in MCF7-LTED cells were accompanied with increases in EP4 protein expression as observed by immunofluorescence (Fig. 1d). Increased EP4 expression in MCF7 LTED cells was also accompanied by increased cAMP in response to the EP4 agonist L-902,688 (Fig. 3b).

### EP4 inactivation decreases estrogen-independent cell growth in MCF7-LTED cells

We sought to determine whether EP4 knockdown would affect the ability of MCF7-LTED cells to grow in the absence of estrogen. EP4 was stably knocked down using lentiviral shRNAs in both MCF7 and MCF7-LTED cells. Visual inspection indicated shEP4 drastically reduced cell proliferation in comparison to a scrambled shRNA control. We further quantified the severe attenuation in growth using reduction in alamarBlue (Supplementary Fig. S3). To better characterize this initial finding, we knocked down mRNA expression using two EP4-targeted siRNAs, and measured relative cellular proliferation seven days after transfection. We verified knockdown at the mRNA level by RT-qPCR (Fig. 2b). Since appropriate antibodies could not be found for Western blot analysis, we validated a decrease in EP4 protein by immunofluorescence (Fig. 2d) and by measurement of the secondary messenger cAMP, which is produced as a result of EP4 activity (Fig. 2c). MCF7-LTED cells treated with siRNAs targeting EP4 mRNA showed a 61% and 59% average reduction in cell proliferation relative to control siRNAs (p < ^10−7^ for each siRNA, Fig. 2a). These siRNAs showed no effect in MCF7 cells, which have negligible levels of EP4 expression at the mRNA and protein levels and thus provide a control for off-target effects.

**Figure 2.**
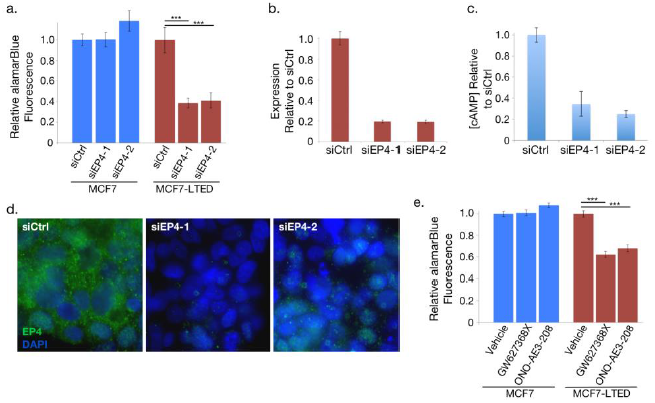
Knockdown or inhibition of EP4 signaling decreases estrogen independent cell proliferation. (a) proliferation of MCF7 cells, which express little EP4, treated with EP4 siRNA and MCF7-LTED cells relative to cells treated with negative control siRNA. (b) RT-qPCR analysis of PTGER4 expression decreases in MCF7-LTED cells treated with two distinct EP4 siRNAs relative to control siRNA. Error bars are standard deviation for three technical replicates. (c) cAMP levels of MCF7-LTED cells treated with EP4 agonist decrease in cells treated with siRNAs to EP4 relative to siRNA controls. (d) Immunofluorescence images of EP4 (green) and DAPI (blue) in MCF7-LTED cells treated with siRNAs targeting EP4 and siRNA controls. Error bars are standard error of the mean (s.e.m.) of three replicates. (e) Cell proliferation of EP4 antagonists in MCF7 and MCF7-LTED cells relative to cells treated with vehicle only. ^***^indicates p < 0.001. Error bars show s.e.m. of three replicates.

We further validated the importance of EP4 for estrogen-independent growth in MCF7-LTED cells using two EP4-specific antagonists that do not inhibit other prostaglandin receptor subtypes: GW627368X and ONO-AE3-208. We observed an average 32% reduction in cellular proliferation for GW627368X-treated and an average 38% reduction for ONO-AE3-208-treated MCF7-LTED cells relative to vehicle alone (p < 10^−7^ for each antagonist, Fig. 2e). Again, MCF7 cells, which have negligible EP4 expression, showed no significant proliferation reduction when treated with antagonists.

### Functional analysis of EP4 signaling in MCF7-LTED cells

EP4 activates two pro-growth signaling pathways: PI3K and PKA; both of which could promote endocrine therapy resistance. MCF7-LTED cells show increased Akt and PI3K activity, and short-term estrogen deprived MCF7 cells are sensitive to the BGT226 and BKM120 PI3K inhibitors^24^. Additionally, EP4 activation can result in phosphorylation and activation of Akt^27^. However, our MCF7-LTED cells are resistant to the effects of the BGT226 and BKM120 PI3K inhibitors^24^. Analysis of Akt and PI3K phosphorylation also showed no changes after treatment of MCF7-LTED cells with EP4 antagonists (Fig. 3a). In addition, we did not find changes in mTOR or MAPK signaling. This suggests that other pathways are likely activated by EP4 in our MCF7-LTED model.

**Figure 3.**
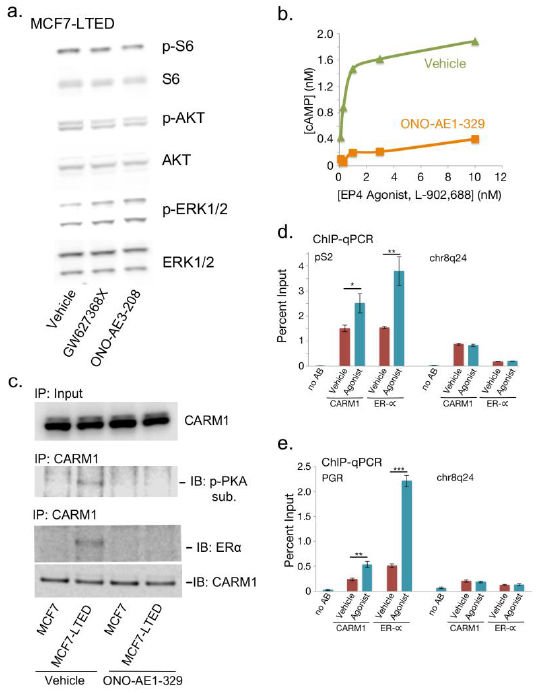
EP4 activates cAMP signaling, and ligand-independent ER activation through CARM1. (a) Western blot analysis of AKT, mTOR (S6 kinase), and MAPK (ERK1/2) pathways in MCF7-LTED cells treated with vehicle or EP4 antagonists. (b) Dose response curve measuring induction of cAMP by EP4 agonist (L-902,688) in MCF7-LTED cells treated with 10nM EP4-antagonist (ONO-AE1-329) or vehicle. (c) Immunoblot of CARM1 in MCF7 and MCF7-LTED cells treated with EP4 antagonist or vehicle in the absence of estrogen. CARM1 immunoprecipitation (IP) followed by western with antibodies for phosphorylated PKA substrate, ERα or CARM1. (d-e) ChIP-qPCR of ERα targets (d) pS2 and (e) PGR with either no antibody (noAB), CARM1 antibody, or ERα antibody in MCF7-LTED cells treated with EP4 agonist or vehicle. Chr8q24 is a gene desert negative control region. * is p-value < 0.05, ** is p-value < 0.01, and *** is p-value < 0.001. Error bars are s.e.m. of three replicates. IB: immunoblot. p-: phosphorylated.

We next hypothesized that *PTGER4* might contribute to endocrine therapy resistance via AC. EP4 activates AC in addition to Akt, and AC activity can induce ERα phosphorylation. Once activated AC produces cAMP as its secondary messenger, and cAMP promotes proliferation through downstream effectors such as PKA^27^. We observed that both the concentration of cAMP in LTED cells and cell proliferation decreased after EP4 reduction by siRNA knockdown (Fig. 2a,Fig. 2c). This suggested that EP4 promotes proliferation in LTED cells via AC and cAMP. To confirm this conclusion, we applied the EP4 specific agonist L-902,688 to MCF7-LTED cells and observed a dramatic increase in cAMP which went away upon inhibition of EP4 with antagonists (Fig. 3b).

A recent report showed that cAMP activates CARM1 via phosphorylation by PKA and drives ligand-independent ER transcriptional activity to promote resistance to tamoxifen^35^. CARM1 is an estrogen-dependent ER co-activator and phosphorylated CARM1 binds directly to ERα^36^. We thus tested whether a similar mechanism caused ligand independent ER signaling in MCF7-LTED cells. CARM1 proteins levels remained unchanged in MCF7-LTED and parental cells in the absence of estrogen even after treatment with EP4 antagonist (Fig. 3c). We found however that CARM1 acted as a PKA substrate in MCF7-LTED cells in the absence of estrogen and this interaction went away after treatment with EP4 antagonist. We further found that ERα showed a strong interaction between CARM1 and ERα in MCF7-LTED cells based on immunoprecipitation of CARM1 followed by immunoblotting with an ERα-specific antibody (Fig. 3c). Treatment with EP4 antagonist removed this interaction, suggesting that the CARM1-ERα interaction is dependent on EP4 signaling. As has been reported previously^36^, in estrogen deprived conditions MCF7 cells showed no activation of CARM1 or association with ERα.

We next used chromatin-immunoprecipitation (ChIP)-qPCR to interrogate the relationship between EP4 and ligand independent binding of ERα in MCF7-LTED cells at two ERα binding sites, the pS2 and PGR promoters. MCF7-LTED cells were treated with agonist or vehicle and subjected to ChIP-qPCR with antibodies to CARM1 or ERα at the pS2 promoter, the PGR promoter, and a negative control gene desert region on chr8q24 (Fig. 3c,Fig. 3d). Both CARM1 and ERα were found to have increased binding upon treatment with EP4-specific agonist relative to cells treated with vehicle. No such gain was observed in the gene desert region. This implies that EP4 signaling can activate ligand-independent ER activation mediated through CARM1.

### ER signaling in MCF7-LTED cells

To understand whether ligand-independent ER signaling is active in MCF7-LTED cells, we compared gene expression in MCF7-LTED cells relative to MCF7 cells that recovered with the addition of estradiol after short-term estrogen deprivation (1 day) using RNA-seq. Using MCF7 cells treated with estradiol we defined a signature of ER signaling based on genes up-regulated two-fold or more. Gene Set Enrichment Analysis (GSEA)^37^ shows that the majority of ER responsive genes are also up-regulated in MCF7-LTED (p < 10^−7^, Supplementary Fig. S6). This is consistent with the finding that our MCF7-LTED cells are responsive to the ER-antagonist fulvestrant^24^ and indicates that ligand-independent ER-signaling is active in MCF7-LTED cells at a genomic level.

### EP4 mRNA expression increased in ER+ breast cancer in the neoadjuvant setting

To assess the potential clinical relevance of our findings, we tested the hypothesis that EP4 expression increases are associated with initial sensitivity to aromatase inhibitor endocrine therapies. Expression profiling was performed for 104 patients before and after neoadjuvant aromatase inhibitor therapy. 74 tumors were determined as ‘aromatase-inhibitor-resistant’ and 30 as ‘aromatase-inhibitor-sensitive’ based on the presence of the proliferation marker Ki67 after treatment (see details in methods). Ki67 is a well-established indicator of clinical outcome in the neoadjuvant setting^38^. We observed that EP4 expression was higher in patients that demonstrated resistance to AI-therapy versus patients that responded to AI-therapy, which is consistent with EP4 expression playing a role in AI-resistance. Interestingly, EP4 expression also significantly increased during neoadjuvant therapy in both responder and non-responder tumor sample. Data from two smaller studies examining mRNA changes during neoadjuvant therapy verify this finding^39,c40^ (Fig. 4). This suggests that in addition to promoting resistance to AIs, EP4 activation may be a response to the loss of ER signaling in ER+ tumors.

**Figure 4.**
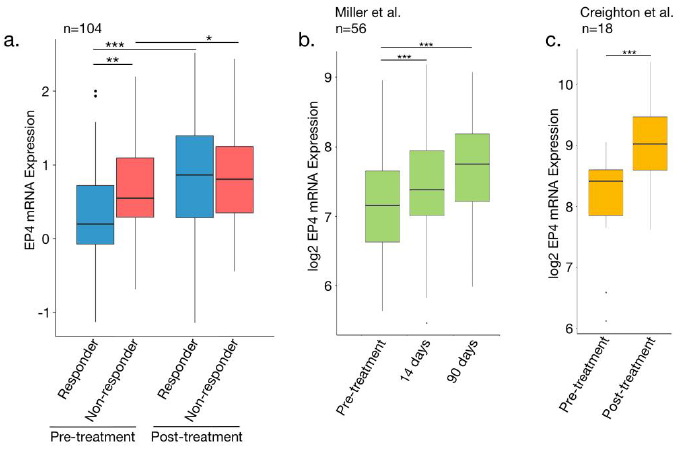
EP4 activates cAMP signaling, and ligand-independent ER activation through CARM1. EP4 expression is higher in patients that fail to respond to neo-adjuvant endocrine therapy. (a) Boxplots of EP4 expression from 104 TCGA tumor samples. Responders and non-responders were defined based on Ki-67 levels after neo-adjuvant therapy. Nonresponders had Ki-67 levels greater than 10%. (b) Expression data from Miller et al. 2011 showing an increase in EP4 expression during pro-longed neo-adjuvant therapy. (c) Expression data from Creighton et al. 2009 for pre-and post-neoadjuvant endocrine therapy. * is p-value < 0.05, ** is p-value < 0.01, and *** is p-value < 0.001. Error bars are s.e.m.

## Discussion

In this study we examined the potential for DNA methylation to facilitate acquired endocrine therapy resistance via EP4 signaling. EP4 signaling also contributes to the proliferation of several cancer types including colon, lung, prostate, ovarian and breast^27^. Antagonists of EP4 have been shown to inhibit metastasis in hormone-resistant murine mammary tumor cells^41^. Further, the contribution of EP4 to endocrine therapy resistance via cell signaling is consistent with its role in the development of castration-resistant prostate cancer via PKA and cAMP^42^.

While our results point to epigenetic regulation of EP4, mutations are unlikely to play a major role in its activation. According to the COSMIC database, EP4 mutations are exceedingly rare in breast cancer, occurring in 0.33% of patients and copy number gains in 1.5%^45^. Sanger sequencing of *PTGER4* in both MCF7 and MC7-LTED cells failed to uncover any mutations that could contribute to the increased activation of EP4. However, analysis of TCGA data indicates that *PTGER4* methylation appears to accompany gene silencing in many breast tumors (Supplementary Fig. S4). Interestingly, *PTGER4* is unmethylated and expressed in normal breast tissue. MCF7 cells treated with demethylating agents increase *PTGER4* mRNA levels, consistent with DNA methylation playing a direct role in its transcriptional regulation^33^. This supports a mechanism whereby an upstream epigenetic change in the EP4 promoter regulates *PTGER4* expression. Namely an increase in DNA methylation that silences *PTGER4* as the tumor forms followed by a decrease in methylation that reactivates the gene to promote resistance to estrogen withdrawal in long-term estrogen deprived cells (Fig. 1).

Increased EP4 levels induce increased cAMP production, which drives ligand-independent activation of ER through the PKA-mediated activation of the co-activator CARM1. Prior work has also indicated that PKA can activate ERα by direct phosphorylation of serines 236 (S236) and 305 (S305). When S236 is phosphorylated, ERα dimerization and DNA binding is still dependent on ligand^36^ and thereby S236 phosphorylation is unlikely to activate ERα in MCF7-LTED cells. Studies in HeLa cells using a phosphomimetic S305E ERα suggested that S305 phosphorylation was sufficient to induce ligand-independent ERα binding to transcriptional targets in the absence of ERα dimerization^43^. However, other studies have instead suggested that S305 phosphorylated ERα is insufficient for ERα activation and that cofactors such as as CARM1^35^ in MCF7 cells and CREB^44^ in CHO were required for ERα binding to DNA and activation of response elements. While we cannot rule out the direct activation of ERα by PKA nor the activation of additional cofactors, our observations suggest that in estrogen deprived conditions EP4 promotes ligand independent activation of ERα through the PKA-mediated activation of the co-activator CARM1.

Another potential mechanism of ligand-independent ER activation is through genetic alterations of the *ESR1* locus. While any individual event is quite rare, as a whole these alterations are becoming an important theme in resistance. Metastatic tumors gain activating *ESR1* ligand binding domain mutations, especially after endocrine therapy^6,46-49^. Translocations of the *ESR1* locus can fuse with activating coding sequence or constitutive promoters that activate *ESR1* in the absence of estrogen^6^. *ESR1* amplification has also been noted in both MCF7-LTED cells^7^ and a xenograft line from a patient tumor resistant to AIs^6^. Rather than contradict our findings, however, this indicates that MCF7-LTED cells remain dependent on ERα signaling. Indeed, MCF7-LTED cells are sensitive to fulvestrant, an ER antagonist that accelerates proteasomal degradation of ER, indicating that these cells remain reliant on ER signaling^24^. GSEA analysis of RNA-seq data from MCF7-LTED cells further shows that MCF7-LTED cells show increased expression of ER target genes. Further, we did not observe a dramatic shift in DNA methylation changes at ER binding sites, which are a surrogate for altered ER binding.

Altogether, these data suggest that increased EP4 expression represents a viable means of developing resistance to aromatase inhibitor therapy. Expression increases in *PTGER4* were associated with decreases in DNA methylation, which is consistent with the idea that DNA methylation has a regulatory role in *PTGER4* expression. Our data indicate that EP4 activation is necessary for estrogen-independent growth. While we cannot completely rule out activation of alternative signaling pathways, our results support the conclusion that EP4 acts through PKA and the co-activator CARM1 to drive ligand-independent ERα activation. EP4 signaling presents a potential therapeutic target for the treatment of AI-resistant breast cancer.

## Materials and Methods

### MCF7 and MCF7-LTED cells

MCF7 cells were from the American Type Culture Collection (Manassas, VA, USA) and maintained in RPMI 1640 (Gibco, Thermo Fisher Scientific, Waltham, MA) supplemented with 5% FBS (Gibco), 10 mM HEPES (Corning, Manassas, VA), 4.5 g/L glucose (Corning), 2 mM L-glutamine (Gibco), 1 mM sodium pyruvate (Corning), and 50 μg/ml gentamicin (Gibco) in a humidified 37°C incubator containing 5% CO_2_. MCF7-LTED cells were previously derived from MCF7 cells^24^ and maintained in the same media except with 5% charcoal stripped FBS (Gibco) and RPMI 1640 without phenol red (Gibco).

### Methyl-MAPS genome-wide methylation analysis

Methyl-MAPS analysis was performed as in Edwards *et al*. with custom barcoded adaptors^28^.
Libraries were made with AB-SOLiD and paired-end sequenced. Sequencing reads were demultiplexed and analyzed using custom perl scripts^28^. Sequencing statistics are in Supplementary Fig. S1.

### RNA-seq

RNA-seq libraries were prepared using NEB Next RNA-seq kit (NEB, Ipswich, MA) with custom barcodes and sequenced with an Illumina HiSeq. Reads were demultiplexed using custom perl scripts. Sequencing statistics are in Supplementary Fig. S1. Reads were mapped to the human genome (hg18) using TopHat (v1.4)^50^. HT-seq was used to assign RefSeq annotations the reads^51^. Statistical analysis was performed using EdgeR^52^. Ontology analysis was performed with Metacore.

### Immunofluorescence

Images were captured on a Zeiss AxioImager Z1 with AxioCam MRc and Axiovision software. The primary antibody was Cayman Chemical Company (Ann Arbor, MI) 101775. The secondary antibody was Jackson ImmunoResearch (West Grove, PA) 111-545-003.

### RT-qPCR

Ep4 mRNA expression was assessed using Applied Biosystems Viia7 with SYBR green. Primer information is in Supplementary Table S1. All qPCR measurements were performed in triplicate.

### Methyl-Screen

Methyl-screen analysis was performed as in Holemon *et al.^34^* with slight modification. Genomic DNA was mock digested, AciI (NEB) digested, McrBC (NEB) digested, or digested with both AciI and McrBC. Real-time quantitative PCR was performed with PTGER4-specific primers: 5’-GCAGCTTTGTCTCTCTTC-3’ and 5’-TACCGAGACCCATGTTG-3’. Unmethylated control gDNA was produced by whole genome amplification of MCF7 gDNA with the REPLI-g kit (Qiagen, Hilden, Germany). Methylated control DNA was produced by treating amplified gDNA with M.SssI (Zymo Research Irvine, CA).

### Cell proliferation

ShRNA targeting EP4 mRNA were from the RNAi core at Washington University in St. Louis. Silencer Select siRNAs were from Ambion (Thermo Fisher Scientific; siCtrl, 4390846; siEP4, 4427037; siEP4-1, ID s60396; siEP4-2, ID s11456). siRNAs were transfected with Lipofectamine RNAiMax (Invitrogen, Thermo Fisher Scientific). When specified, EP4 antagonist GW627368X was used at 3.3 μM and ONO-AE3-208 was used at 10 μM. AlamarBlue assays were performed at 7 days for MCF7-LTED and 4 days for MCF7 cells. One-tenth volume of alamarBlue reagent (AbD Serotec BUF012, Bio-Rad, Oxford, UK) was added to each replicate. Eight replicates were performed for MCF7-LTED cells and 4 replicates for MCF7 cells. p-values were computed using a one-way ANOVA followed post-hoc by Tukey’s HSD (honest significant difference) test. Bartlett’s test of the homogeneity of variances was insignificant (alpha = 0.05) for all comparisons under the null-hypothesis of unequal variances.

### ChlP-qPCR

Cells were grown in 10 cm dishes to 85% confluence. The cells were starved, by exchanging growth media for reduced-serum media containing 0.5% charcoal stripped FBS, for 18 hours before activation of EP4 with 10nM L902,688 EP4 agonist (Cayman Chemical) for 2 hours. An ethanol-treated sample served as a vehicle control. Afterward, cross-linking was initiated by drop-wise addition of 37% formaldehyde to a final concentration of 1% at room temperature. Cross-linking ran for 10 minutes and was quenched for 5 minutes at room temperature by adding 0.5ml of 2.5M glycine. Quenched cells were washed twice with 10ml ice-cold PBS, removed from the plate by scraping in ice-cold DPBS (Gibco) with protease inhibitors, and pelleted. Cell pellets were resuspended in 350 ul ChIP lysis buffer (1% SDS, 10 mM EDTA, 50 mM Tris-HCl pH 8.0) with protease inhibitors. Lysed cells were sonicated and 100 ug of protein were brought to a final volume of 50ul lysis buffer with protease inhibitors. 9 ul of CARM1 antibody (CST, (3H2) Mouse mAb #12495) or 2ul ER antibody (Santa Cruz sc-7207X) was added and incubated overnight at 4°C.

Antibody bound proteins were purified by adding samples to 25μl Protein A/G magnetic beads (Pierce) that were prewashed twice with ChIP Dilution Buffer. The samples were incubated with the beads 2 hours at 4 °C. Afterward the beads were washed with 0.5 ml of each of the following ice-cold buffers: 1) low salt immune complex wash buffer (EMD-Millipore 20-154), one wash; 2) high salt immune complex wash buffer (EMD-Millipore 20-155), one wash; 3) LiCl immune complex wash buffer (EMD-Millipore 20-156), one wash; 4) 1X TE-T, two washes (10mM TrisHCl, 1mM EDTA, 0.1% Triton, pH 8). Elution buffer (1% SDS + 0.1M NaHCO3) was prepared fresh from 10X stocks. 200 μl room temperature elution buffer was added, and the samples were incubated at room temperature for 30 min to elute the protein. Afterward, 1 μl of 10 mg/mL RNase A was added and the samples were incubated overnight at 65°C. The Qiagen PCR Purification kit was used to purify the DNA in a final volume of 60μl Buffer EB.

Real-time quantitative PCR was performed on an Applied Biosystems Viia7 using SYBR green. Primers are in Supplementary Table S2. Values are reported as percent relative to input. All experiments were performed in triplicate and p-values are computed using Student’s t-test.

### cAMP measurements

CAMP measurements were made using the Cyclic AMP XP^®^ Assay Kit (Cell Signaling Technologies) according to the manufacturer’s instructions. MCF7-LTED cells were washed twice with warm PBS and then pre-treated with 0.5 mM IBMX in serum-free medium for 30 min prior to addition of L-902,688, at the indicated concentrations, or 1 μM forskolin. For combined antagonist/agonist treatments, cells were pre-treated 10 min with ONO-AE3-208 before addition of L-902,688. For EP4 knock down experiments, cAMP measurements were made 2 days post transfection. cAMP measurements were quantitated as follows: %Activity = 100 × [(A–A_basal_)/(A_max_–A_basal_)], where A is the absorbance of the agonist treated sample, A_max_ is the absorbance of the forskolin treated sample, and A_basal_ is the absorbance of the vehicle treated sample.

### CARM1-IP - ER blot

MCF7-LTED cells were grown to confluence in 10 cm dishes. Before harvesting, cells were grown in media with low serum concentration (0.5%). The next day, the cells were incubated with 10 mL of RPMI (no serum) containing 0.5 mM IBMX for 30 minutes. The cells were then treated with 10 nM L-902,688 in 10 mL of RPMI + 0.5 mM IBMX and harvested after 0, 5, 15, or 30 minutes. After washing with ice-cold PBS, cells were lysed with 1 mL of RIPA buffer (1 M tris pH 8.0, 5 M NaCl, 1% NP-40, 0.5% sodium deoxycholate, 0.1% SDS, and 2 mM EDTA with freshly added PMSF to 1 mM final and 1X Halt protease and phosphatase inhibitor cocktail [ThermoFisher Scientific]). Cells were scraped loose, the cell lysate was transferred to a microcentrifuge tube, and the tube was put on a rotator for 30 minutes at 4°C. The supernatant was isolated by centrifugation at 12,000xg for 20 minutes at 4°C.

Immunoprecipitation was performed by addition of mouse anti-CARM1 (Cell Signaling Technology #12495) and precipitation overnight at 4°C. Precipitated proteins were collected with the Dynabeads Protein G magnetic beads. 25 uL of beads were washed with 500 uL of RIPA buffer before adding the immunoprecipitated proteins. This was then incubated at 4 °C for 3 hours. The protein-bound beads were then washed three times with 0.5 mL RIPA with 10 mM NaF and 1 mM sodium pyrophosphate and resuspended in 0.5 mL of wash buffer. The protein was eluted in 40 uL of 2x NuPAGE LDS sample buffer at 90°C for 10 minutes. Final analysis of immunoprecipitated proteins occurred by Western blot on 4-12% gradient PAGE gels (Life Tech.). ERα was identified using a rabbit polyclonal antibody (Santa Cruz Biotech sc-7207) to minimize signal from the mouse anti-CARM1 IgG used for immunoprecipitation. The blot was stripped using Restore™ Western Blot Stripping Buffer (ThermoFisher), and reblotted with rabbit anti-CARM1 antibody (Cell Signaling Technologies #3379). Alexa Fluor^®^ 680 or Alexa Fluor^®^ 790 labeled secondary antibodies were from Jackson ImmunoResearch. Blots were imaged using the Odyssey^®^ Fc (LI-COR Biosciences).

### The Cancer Genome Atlas data analysis

Infinium 450k methylation, RNA-seq expression and clinical data were downloaded from the The Cancer Genome Atlas (TCGA) data portal. P-values were computed using Mann-Whitney U.

### Expression Analysis of ER+ breast cancer in neoadjuvant setting

Expression analysis was performed for patients enrolled in the POL^53^ and ACOSOG (American College of Surgeons Oncology Group) Z1031^54^ trials. Postmenopausal women with ER+ Stage II/III breast cancer were treated with letrozole, anastrozole, or exemestane for 16-18 weeks. Biopsies were performed before and after treatment. Patients were labeled as resistant if 10% or more of malignant cells stained positive for the proliferative marker Ki67 at 16 weeks. Expression analysis was performed using Agilent Human Gene Expression 4x44K v2 Microarray. P-values for *PTGER4* expression were computed using Mann-Whitney U.

## Acknowledgements

We thank C. De Guzman-Strong and S. Matkovich for their comments on the manuscript. This work was supported by grants NIH R00 CA127360, NIH R21 LM011199, Department of Defense W81XWH-11-1-0401 and a Siteman Cancer Center Breast Cancer Program career development award to J.R.E. We also thank the Genome Technology Access Center at the Washington University School of Medicine for help with genomic analysis (partially supported by NIH P30 CA91842 and NIH UL1 TR000448).

## References

1 Harvey JM, Clark GM, Osborne CK, Allred DC. Estrogen Receptor Status by Immunohistochemistry Is Superior to the Ligand-Binding Assay for Predicting Response to Adjuvant Endocrine Therapy in Breast Cancer. Journal of Clinical Oncology 1999; 17: 1474–1474.

2 Johnston SRD, Dowsett M. Aromatase inhibitors for breast cancer: lessons from the laboratory. Nat Rev Cancer 2003; 3: 821–831.

3 Musgrove EA, Sutherland RL. Biological determinants of endocrine resistance in breast cancer. Nat Rev Cancer 2009; 9: 631–643.

4 Dowsett M, Cuzick J, Ingle J, Coates A, Forbes J, Bliss J et al. Meta-analysis of breast cancer outcomes in adjuvant trials of aromatase inhibitors versus tamoxifen. J Clin Oncol 2010; 28: 509–518.

5 Ma CX, Reinert T, Chmielewska I, Ellis MJ. Mechanisms of aromatase inhibitor resistance. Nat Rev Cancer 2015; 15: 261–275.

6 Li S, Shen D, Shao J, Crowder R, Liu W, Prat A et al. Endocrine-Therapy-Resistant ESR1 Variants Revealed by Genomic Characterization of Breast-Cancer-Derived Xenografts. Cell reports 2013; 4: 1116–1130.

7 Aguilar H, Solé X, Bonifaci N, Serra-Musach J, Islam A, López-Bigas N et al. Biological reprogramming in acquired resistance to endocrine therapy of breast cancer. Oncogene 2010. doi:10.1038/onc.2010.333.

8 Zilli M, Grassadonia A, Tinari N, Di Giacobbe A, Gildetti S, Giampietro J et al. Molecular mechanisms of endocrine resistance and their implication in the therapy of breast cancer. 2009; 1795: 62–81.

9 Ellis MJ, Ding L, Shen D, Luo J, Suman VJ, Wallis JW et al. Whole-genome analysis informs breast cancer response to aromatase inhibition. Nature 2012; 486: 353–360.

10 Iorns E, Turner NC, Elliott R, Syed N, Garrone O, Gasco M et al. Identification of CDK10 as an important determinant of resistance to endocrine therapy for breast cancer. Cancer Cell 2008; 13: 91–104.

11 Kato S, Endoh H, Masuhiro Y, Kitamoto T, Uchiyama S, Sasaki H et al. Activation of the estrogen receptor through phosphorylation by mitogen-activated protein kinase. Science (New York, NY) 1995; 270: 1491–1494.

12 Le Goff P, Montano MM, Schodin DJ, Katzenellenbogen BS. Phosphorylation of the human estrogen receptor. Identification of hormone-regulated sites and examination of their influence on transcriptional activity. J Biol Chem 1994; 269: 4458–4466.

13 Feinberg AP, Ohlsson R, Henikoff S. The epigenetic progenitor origin of human cancer.

14 Kulis M, Esteller M. 2 - DNA Methylation and Cancer. In: Ushijima ZHAT (ed). Advances in Genetics. Academic Press, 2010, pp 27–56.

15 Busslinger M, Hurst J, Flavell RA. DNA methylation and the regulation of globin gene expression. Cell 1983; 34: 197–206.

16 Ottaviano YL, Issa J-P, Parl FF, Smith HS, Baylin SB, Davidson NE. Methylation of the Estrogen Receptor Gene CpG Island Marks Loss of Estrogen Receptor Expression in Human Breast Cancer Cells. Cancer Res 1994; 54: 2552–2555.

17 Ferguson AT, Lapidus RG, Baylin SB, Davidson NE. Demethylation of the Estrogen Receptor Gene in Estrogen Receptor-negative Breast Cancer Cells Can Reactivate Estrogen Receptor Gene Expression. Cancer Res 1995; 55: 2279–2283.

18 Yang X, Phillips DL, Ferguson AT, Nelson WG, Herman JG, Davidson NE. Synergistic activation of functional estrogen receptor (ER)-alpha by DNA methyltransferase and histone deacetylase inhibition in human ER-alpha-negative breast cancer cells. Cancer Res 2001; 61:7025–7029.

19 Yang X, Ferguson AT, Nass SJ, Phillips DL, Butash KA, Wang SM et al. Transcriptional activation of estrogen receptor alpha in human breast cancer cells by histone deacetylase inhibition. Cancer Res 2000; 60: 6890–6894.

20 Sabnis GJ, Goloubeva O, Chumsri S, Nguyen N, Sukumar S, Brodie AMH. Functional activation of the estrogen receptor-α and aromatase by the HDAC inhibitor entinostat sensitizes ER-negative tumors to letrozole. Cancer Res 2011; 71: 1893–1903.

21 Keen JC, Yan L, Mack KM, Pettit C, Smith D, Sharma D et al. A novel histone deacetylase inhibitor, scriptaid, enhances expression of functional estrogen receptor alpha (ER) in ER negative human breast cancer cells in combination with 5-aza 2′-deoxycytidine. Breast Cancer Res Treat 2003; 81: 177–186.

22 Giacinti L, Giacinti C, Gabellini C, Rizzuto E, Lopez M, Giordano A. Scriptaid effects on breast cancer cell lines. Journal of cellular physiology 2012; 227: 3426–3433.

23 Zhou Q, Shaw PG, Davidson NE. Inhibition of histone deacetylase suppresses EGF signaling pathways by destabilizing EGFR mRNA in ER-negative human breast cancer cells. Breast Cancer Res Treat 2008; 117: 443–451.

24 Sanchez CG, Ma CX, Crowder RJ, Guintoli T, Phommaly C, Gao F et al. Preclinical modeling of combined phosphatidylinositol-3-kinase inhibition with endocrine therapy for estrogen receptor-positive breast cancer. Breast Cancer Res 2011; 13: R21.

25 Crowder RJ, Phommaly C, Tao Y, Hoog J, Luo J, Perou CM et al. PIK3CA and PIK3CB inhibition produce synthetic lethality when combined with estrogen deprivation in estrogen receptor-positive breast cancer. Cancer Res 2009; 69: 3955–3962.

26 Chan CMW, Martin L-A, Johnston SRD, Ali S, Dowsett M. Molecular changes associated with the acquisition of oestrogen hypersensitivity in MCF-7 breast cancer cells on long-term oestrogen deprivation. J Steroid Biochem Mol Biol 2002; 81: 333–341.

27 Yokoyama U, Iwatsubo K, Umemura M, Fujita T, Ishikawa Y. The prostanoid EP4 receptor and its signaling pathway. Pharmacological Reviews 2013; 65: 1010–1052.

28 Edwards JR, O'Donnell AH, Rollins RA, Peckham HE, Lee C, Milekic MH et al. Chromatin and sequence features that define the fine and gross structure of genomic methylation patterns. Genome Res 2010; 20: 972–980.

29 van Agthoven T, van Agthoven TL, Dekker A, Foekens JA, Dorssers LC. Induction of estrogen independence of ZR-75-1 human breast cancer cells by epigenetic alterations. Mol Endocrinol 1994; 8: 1474–1483.

30 Vanderkraats ND, Hiken JF, Decker KF, Edwards JR. Discovering high-resolution patterns of differential DNA methylation that correlate with gene expression changes. Nucleic Acids Res 2013; 41: 6816–6827.

31 Irizarry RA, Ladd-Acosta C, Wen B, Wu Z, Montano C, Onyango P et al. The human colon cancer methylome shows similar hypo-and hypermethylation at conserved tissue-specific CpG island shores. Nat Genet 2009; 41: 178–186.

32 Fritah A, Saucier C, De Wever O, Bracke M, Bieche I, Lidereau R et al. Role of WISP-2/CCN5 in the maintenance of a differentiated and noninvasive phenotype in human breast cancer cells. Mol Cell Biol 2008; 28: 1114–1123.

33 Kim JH, Kang S, Kim TW, Yin L, Liu R, Kim SJ. Expression profiling after induction of demethylation in MCF-7 breast cancer cells identifies involvement of TNF-α mediated cancer pathways. Molecules and cells 2012; 33: 127–133.

34 Holemon H, Korshunova Y, Ordway JM, Bedell JA, Citek RW, Lakey N et al. MethylScreen: DNA methylation density monitoring using quantitative PCR. BioTechniques 2007; 43: 683–693.

35 Carascossa S, Dudek P, Cenni B, Briand P-A, Picard D. CARM1 mediates the ligand-independent and tamoxifen-resistant activation of the estrogen receptor alpha by cAMP. 2010; 24: 708–719.

36 Chen D, Pace PE, Coombes RC, Ali S. Phosphorylation of human estrogen receptor alpha by protein kinase A regulates dimerization. Mol Cell Biol 1999; 19: 1002–1015.

37 Subramanian A, Tamayo P, Mootha VK, Mukherjee S, Ebert BL, Gillette MA et al. Gene set enrichment analysis: a knowledge-based approach for interpreting genome-wide expression profiles. Proceedings of the National Academy of Sciences of the United States of America 2005; 102: 15545–15550.

38 Ellis MJ, Tao Y, Luo J, A’Hern R, Evans DB, Bhatnagar AS et al. Outcome Prediction for Estrogen Receptor-Positive Breast Cancer Based on Postneoadjuvant Endocrine therapy Tumor Characteristics. J Natl Cancer Inst 2008; 100: 1380–1388.

39 Miller TW, Balko JM, Ghazoui Z, Dunbier A, Anderson H, Dowsett M et al. A gene expression signature from human breast cancer cells with acquired hormone independence identifies MYC as a mediator of antiestrogen resistance. Clin Cancer Res 2011; 17: 2024–2034.

40 Creighton CJ, Li X, Landis M, Dixon JM, Neumeister VM, Sjolund A et al. Residual breast cancers after conventional therapy display mesenchymal as well as tumor-initiating features. Proceedings of the National Academy of Sciences of the United States of America 2009; 106:13820–13825.

41 Ma X, Kundu N, Rifat S, Walser T, Fulton AM. Prostaglandin E receptor EP4 antagonism inhibits breast cancer metastasis. Cancer Res 2006; 66: 2923–2927.

42 Terada N, Shimizu Y, Kamba T, Inoue T, Maeno A, Kobayashi T et al. Identification of EP4 as a potential target for the treatment of castration-resistant prostate cancer using a novel xenograft model. Cancer Res 2010; 70: 1606–1615.

43 Tharakan R, Lepont P, Singleton D, Kumar R, Khan S. Phosphorylation of estrogen receptor alpha, serine residue 305 enhances activity. Molecular and Cellular Endocrinology 2008; 295: 70–78.

44 Lazennec G, Thomas JA, Katzenellenbogen BS. Involvement of cyclic AMP response element binding protein (CREB) and estrogen receptor phosphorylation in the synergistic activation of the estrogen receptor by estradiol and protein kinase activators. J Steroid Biochem Mol Biol 2001; 77: 193–203.

45 Forbes SA, Beare D, Gunasekaran P, Leung K, Bindal N, Boutselakis H et al. COSMIC: exploring the world’s knowledge of somatic mutations in human cancer. Nucleic Acids Res 2015; 43: D805–11.

46 Zhang QX, Borg A, Wolf DM, Oesterreich S, Fuqua SA. An estrogen receptor mutant with strong hormone-independent activity from a metastatic breast cancer. Cancer Res 1997; 57: 1244–1249.

47 Toy W, Shen Y, Won H, Green B, Sakr RA, Will M et al. ESR1 ligand-binding domain mutations in hormone-resistant breast cancer. Nat Genet 2013; 45: 1439–1445.

48 Merenbakh-Lamin K, Ben-Baruch N, Yeheskel A, Dvir A, Soussan-Gutman L, Jeselsohn R et al. D538G mutation in estrogen receptor-α: A novel mechanism for acquired endocrine resistance in breast cancer. Cancer Res 2013; 73: 6856–6864.

49 Robinson DR, Wu Y-M, Vats P, Su F, Lonigro RJ, Cao X et al. Activating ESR1 mutations in hormone-resistant metastatic breast cancer. Nat Genet 2013; 45: 1446–1451.

50 Trapnell C, Pachter L, Salzberg SL. TopHat: discovering splice junctions with RNA-Seq. Bioinformatics 2009; 25: 1105–1111.

51 Anders S, Pyl PT, Huber W. HTSeq--a Python framework to work with high-throughput sequencing data. Bioinformatics 2015 31: 166–169.

52 Robinson MD, McCarthy DJ, Smyth GK. edgeR: a Bioconductor package for differential expression analysis of digital gene expression data. Bioinformatics 2010 26: 139–140.

53 Olson JAJr, Budd GT, Carey LA, Harris LA, Esserman LJ, Fleming GF et al. Improved surgical outcomes for breast cancer patients receiving neoadjuvant aromatase inhibitor therapy: results from a multicenter phase II trial. - PubMed - NCBI. Journal of the American College of Surgeons 2009; 208: 906–914.

54 Ellis MJ, Suman VJ, Hoog J, Lin L, Snider J, Prat A et al. Randomized phase II neoadjuvant comparison between letrozole, anastrozole, and exemestane for postmenopausal women with estrogen receptor-rich stage 2 to 3 breast cancer: clinical and biomarker outcomes and predictive value of the baseline PAM50-based intrinsic subtype--ACOSOG Z1031. J Clin Oncol 2011 29: 2342–2349.

